# Mixtures that Matter: Correlation Patterns in Antibacterial and Cytotoxic Activities of Five Hop Isolates

**DOI:** 10.64898/2026.04.13.718148

**Authors:** Luisa Kober, Luca von Karger, Kathrin Castiglione

## Abstract

Antimicrobial resistance (AMR) has resulted in the need for the development of alternative strategies for combating pathogens and growth promotion of poultry, including the use of plant-derived compounds such as hop (*Humulus lupulus*) isolates. The present study evaluates the correlation patterns of the biological activity of five major hop isolates (humulone, lupulone, isohumulone, xanthohumol, and isoxanthohumol) against *Bacillus subtilis, Micrococcus luteus*, and a chicken cell line UMNSAH/DF-1 using a two-dimensional checkerboard assay. Fractional inhibitory concentrations (∑FIC) were used to classify interactions as additive, synergistic, or antagonistic, and selectivity indices assessed antibacterial versus cytotoxic effects. On *B. subtilis*, combinations were predominantly additive, whereas *M. luteus*, in contrast, showed variable interactions, including also synergistic (humulone + lupulone) and antagonistic combinations (isohumulone + isoxanthohumol), demonstrating the impact of the metabolic resilience of the target organism. Cytotoxicity in UMNSAH/DF-1 cells was largely additive, with synergistic effects observed only for isomerized compounds. Selectivity analysis highlighted humulone-lupulone combinations as most favorable, offering high antibacterial activity with minimal cytotoxicity. These results provide novel insights for selecting hop isolate combinations for the development of phytogenic feed additives (PFAs), emphasizing that both compound composition and target organism physiology critically shape efficacy and safety outcomes.

## 1. INTRODUCTION

Antimicrobial resistance (AMR) in bacterial pathogens is a global challenge associated with high morbidity and mortality. The presence of multidrug resistant patterns in bacteria has led to the emergence of infections that are difficult to treat, and in some cases, even untreatable, with conventional antimicrobials [1]. According to the One Health programme of the United Nations, by the year 2050, AMR will have become the foremost cause of mortality, along with cancer, with both accounting for 10,000,000 deaths worldwide [2]. The administration of subtherapeutic or prophylactic doses of antibiotics remains prevalent, particularly in the field of livestock farming of poultry, despite the existence of regions and countries that have already instituted prohibitions on this practice. To counteract this, research is ongoing into alternative solutions, many of which are plant-based, such as hops (*Humulus lupulus*), which have been shown to possess antibacterial properties [3].

Plant-derived products and extracts are challenging to research due to their chemical complexity. In comparison with single-compound substances, plant materials frequently contain a variety of bioactives, including primary and secondary metabolites, whose composition can differ depending on species, growth conditions, and extraction methods. This heterogeneity complicates the attribution of observed biological effects to specific compounds [4]. Nevertheless, plants offer the advantage of being a renewable and sustainable source of bioactive compounds [5]. Given the wide range of substances present in hops, the focus should be on those that constitute the largest bioactive proportion. These include humulone (alpha-acids) and lupulone (beta-acids), which are well-known for their antibacterial properties. Xanthohumol is another potential antimicrobial that can be added to this list. The two isoforms isohumulone and isoxanthohumol, which are produced at high temperatures, e.g. during brewing, are also of interest, as these compounds are major drivers of the microbiological stability of beer and also have altered solubility in polar media, which could be advantageous for certain applications [6].

In order to better interpret the interaction of these hop isolates, it is essential to understand the various possibilities for compound correlation. When two or more active ingredients are used together, three different effects can occur: additivity, synergism, or antagonism [7]. The effects can be described as follows: additive interactions, in which the response to a combination of drugs corresponds exactly to the dose-response relationship of the individual drugs; synergistic interactions, in which the response is greater than expected; and antagonistic interactions, in which the response is less than expected [8]. **Fig.*1*** shows the underlying mechanisms for synergistic and antagonistic behaviour.

**Fig. 1.**
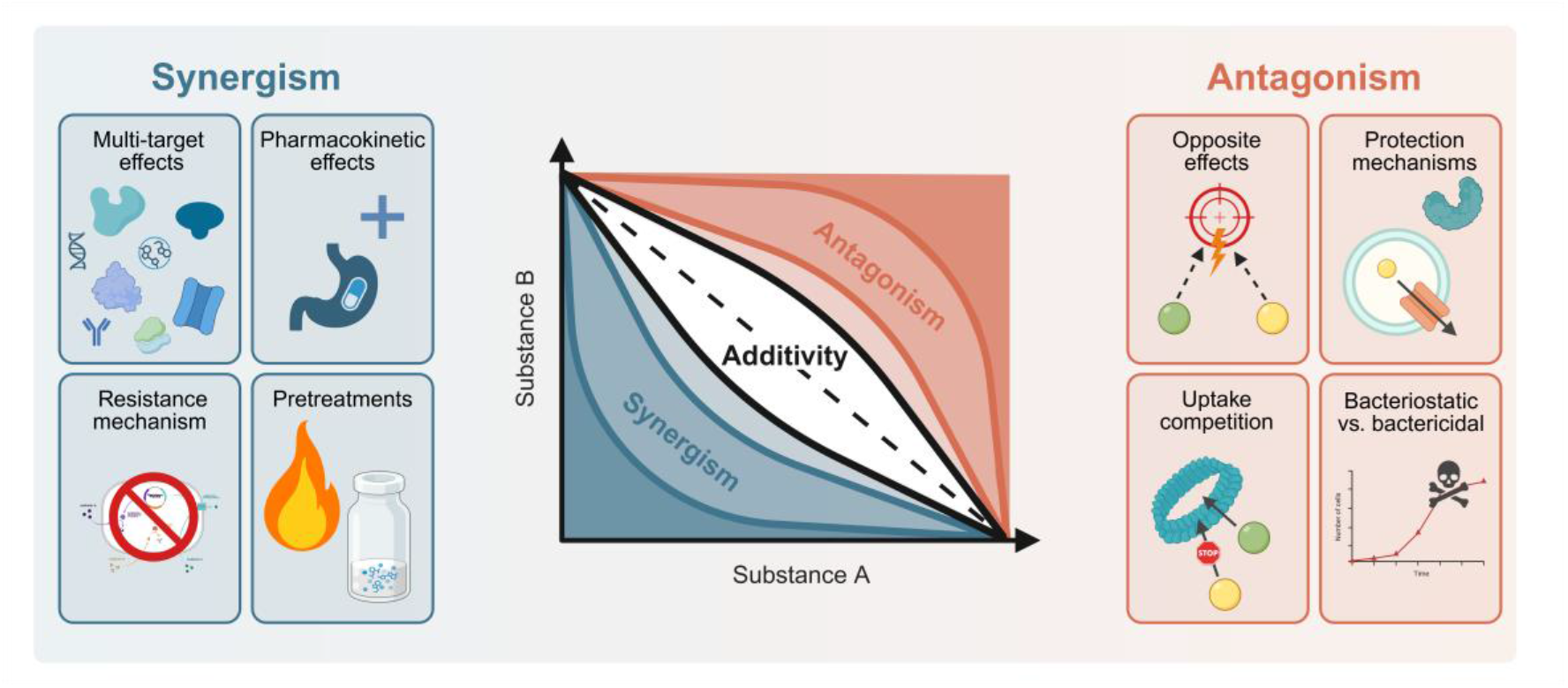
Schematic overview of correlation mechanisms than can occur leading to synergism or antagonism. The diagram shows an isobologram in which the dotted line represents the additive relationship between two substances. All points significantly below this line (blue area) symbolize synergism, while all points significantly above the line (red area) symbolize antagonism [9-13]. This illustration is created with BioRender.

Synergistic effects between antimicrobial agents can be categorised into four principal mechanisms. First, multi-target effects arise when compounds act simultaneously on distinct targets such as enzymes, substrates, metabolites, receptors, transport proteins, DNA/RNA, ribosomes, antibodies, physicochemical reactions, or even signalling cascades. Second, pharmacokinetic or physicochemical effects describe situations in which compounds without specific pharmacological activity enhance the solubility and absorption of active drugs, thereby improving their overall bioavailability. Third, interactions with resistance mechanisms play a crucial role, for example, when enzymes that would otherwise inactivate antibiotics are inhibited, or when efflux pumps responsible for exporting antibiotics out of bacterial cells are blocked. Finally, elimination or neutralization of undesired reactions does not represent a form of synergy in a strict pharmacological sense, but rather involves pretreatment or modification of agents, such as heat inactivation or the addition of other substances, to reduce adverse interactions and facilitate therapeutic efficacy [9].

Furthermore, antagonistic interactions between antimicrobial agents can also be explained. One mechanism involves competition for identical or overlapping targets, whereby two compounds interfere with each other’s binding or functional activity at the same molecular site, thereby reducing overall efficacy [10]. Second, the induction of bacterial stress responses or protective mechanisms can result in the diminished activity of a second drug. For instance, the activation of efflux pumps, changes in membrane permeability or the expression of protective enzymes may be triggered by one compound, hindering the other [11]. Third, uptake can be hindered when one antibiotic alters the absorption, distribution or stability of the other [12]. Finally, one cause is the combination of bacteriostatic and bactericidal agents, whereby growth inhibition by the former prevents the other from exerting its activity on actively dividing cells [13].

The interaction between hop isolates is, unfortunately, not yet well understood and there is a lack of studies comparing individual hop isolates in the literature. However, this information is crucial for understanding how hop products could work as a plant-based alternative (phytogenic feed additive, PFA) to antibiotic growth promotors (AGPs). AGPs are compounds that are not primarily used to treat diseases, but rather to promote growth and efficiency in animal production. They alter intestinal flora, reducing the occurrence of certain pathogenic bacteria and improving nutrient absorption [14].

After evaluating the effects of individual hop compounds [15], it is necessary to take the next steps in the direction of whole hop products. Therefore, we investigated the interaction effects of the hop isolates humulone, lupulone, isohumulone, xanthohumol and isoxanthohumol on two bacterial strains and a chicken cell line with a two-dimensional checkerboard assay. Based on these results, we can conclude which hop compounds should be enriched or depleted in hop extracts to produce a sustainable poultry farming product based on hops as a raw and renewable material.

## 2. RESULTS

### 2.1. Enhancing solubility of hop isolates in aqueous media through β-cyclodextrin

Due to the hydrophobic character of the hop isolates analysed here, a complete dissolution in aqueous medium cannot be assumed. This was demonstrated by the visually perceptible precipitation of the pure substances when the ethanolic stock solutions were diluted in aqueous medium. The Checkerboard assay requires rather high concentrations of the respective dissoluted hop isolates, as a 4-fold dilution occurs throughout the assay. Due to the comparatively poor solubility in bacterial media NB and TSB, 10 mM β-cyclodextrin was added to all isolate preparations that were tested for antibacterial activity. Preliminary experiments (**Fig.*2***) showed that the antibacterial effect of hop isolates was not altered by the addition of CD (+ CD), suggesting that CD does not bind the substances too tightly in its hydrophobic cavity and that the molecules easily distribute between their free and bound form [16].

**Fig. 2.**
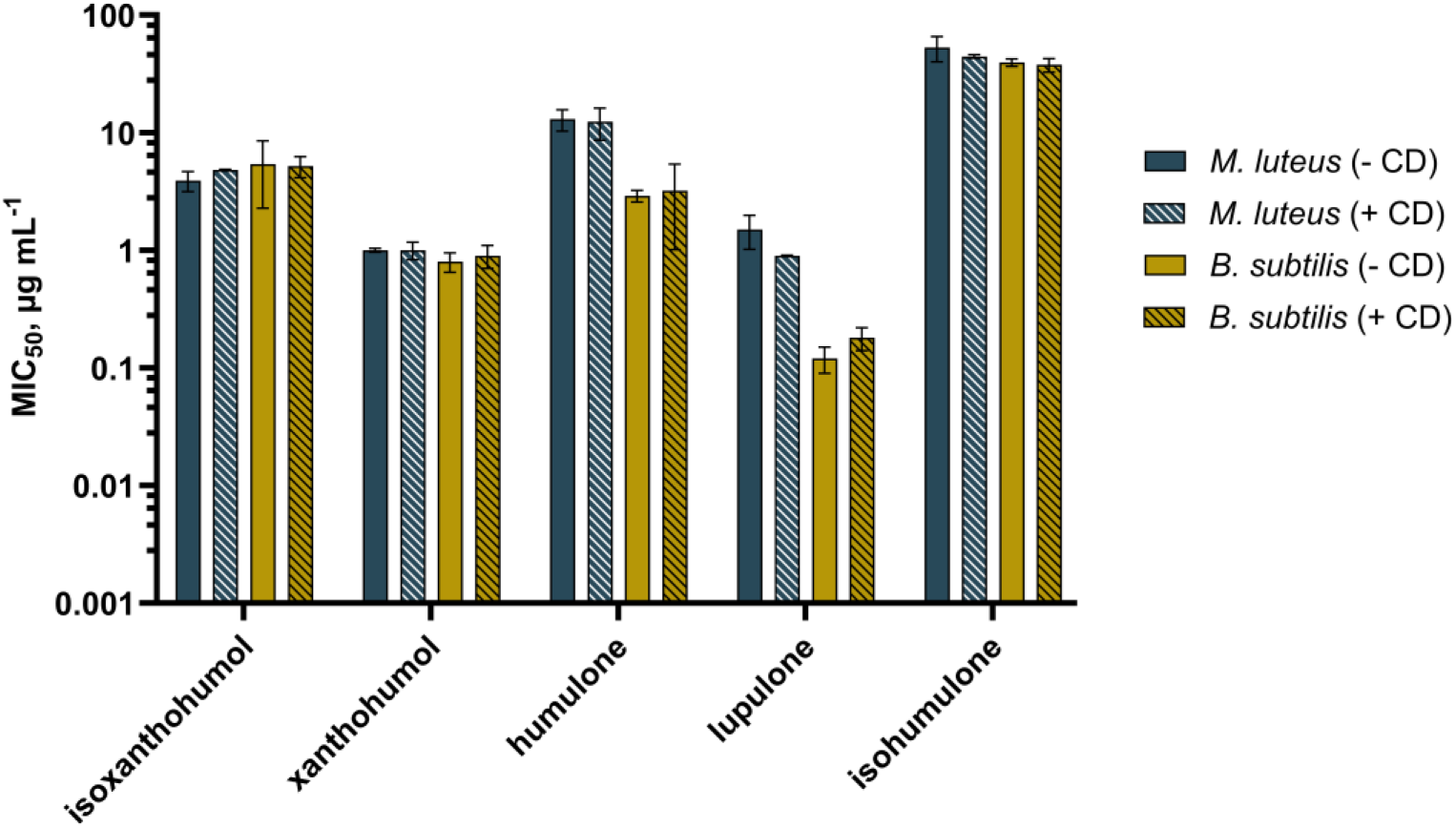
Influence of ß-cyclodextrin (CD) on the antibacterial activity of five hop isolates isoxanthohumol, xanthohumol, humulone, lupulone, and isohumulone against *Micrococcus luteus* (*M. luteus*) and *Bacillus subtilis* (*B. subtilis*). MIC_50_ are biological duplicates.

Furthermore, it can be assumed that the effect of CD would also be minimized when evaluating the data, since a potential influence would be eliminated mathematically (see Eq. (1)).

**Fig.*3*** shows the different solubilities of the five hop isolates after addition of 10 mM ß-cyclodextrin.

**Fig. 3.**
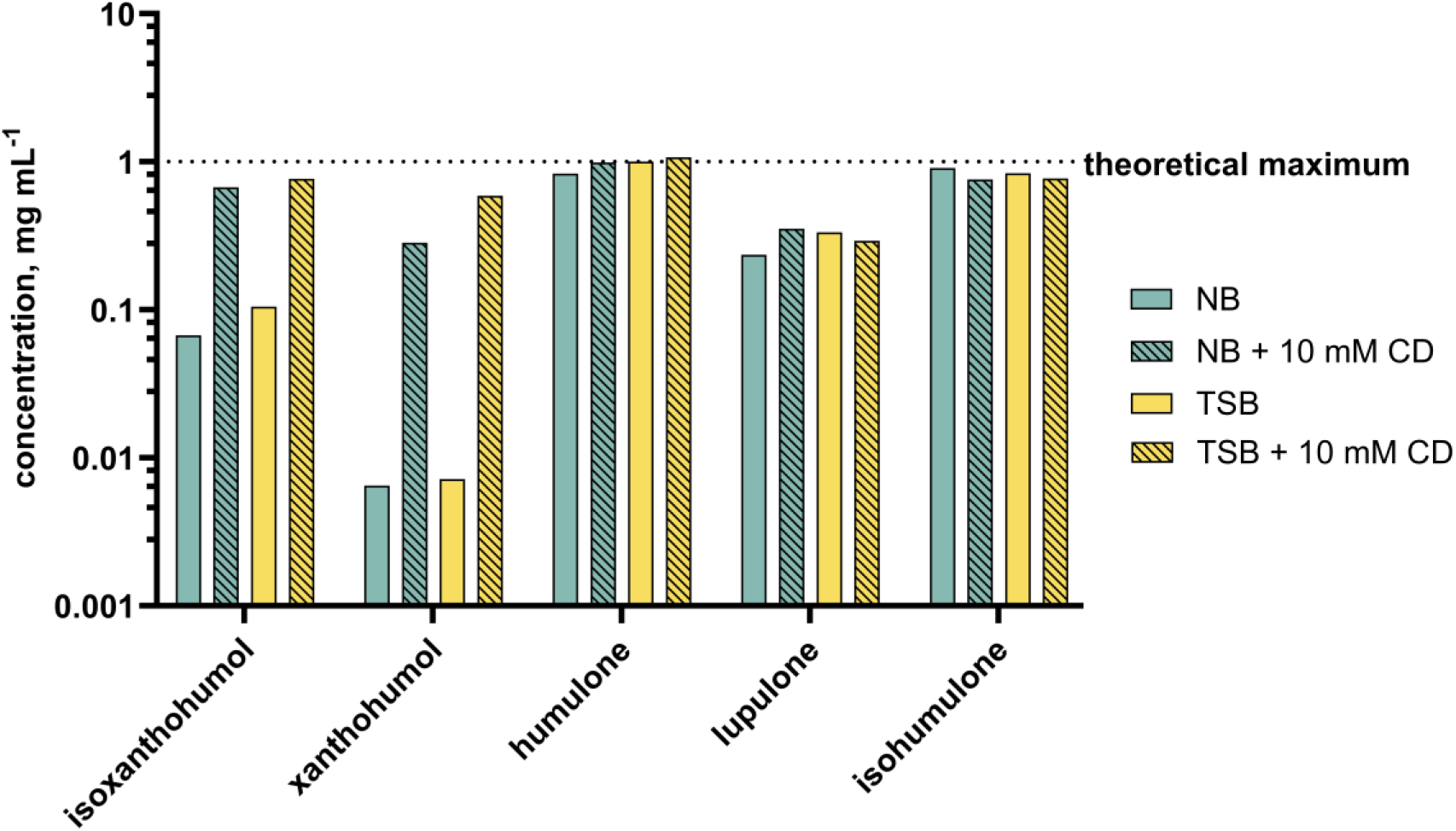
Solubility of five hop isolates isoxanthohumol, xanthohumol, humulone, lupulone, and isohumulone in two bacterial media NB and TSB each with 10% (v/v) ethanol content, with and without the addition of 10 mM β-cyclodextrin (CD).

For bitter acids humulone, lupulone, and isohumulone, almost no change in solubility was detected. A clear improvement in solubility was observed for isoxanthohumol and xanthohumol. Isoxanthohumol concentration in NB was increased by a factor of 10, and in TSB by a factor of 7. Xanthohumol concentrations could even be enhanced by a factor of 44 (NB) and 82 (TSB), which enabled the checkerboard assay to be performed.

### 2.2. Antibacterial correlation patterns against *Micrococcus luteus* and *Bacillus subtilis*

After subtracting the background from the signal values, a decreasing value gradient emerged from the lower left well (growth control) to the upper right well. This is illustrated exemplary in **Fig.*4*** in form of a heat map.

**Fig. 4.**
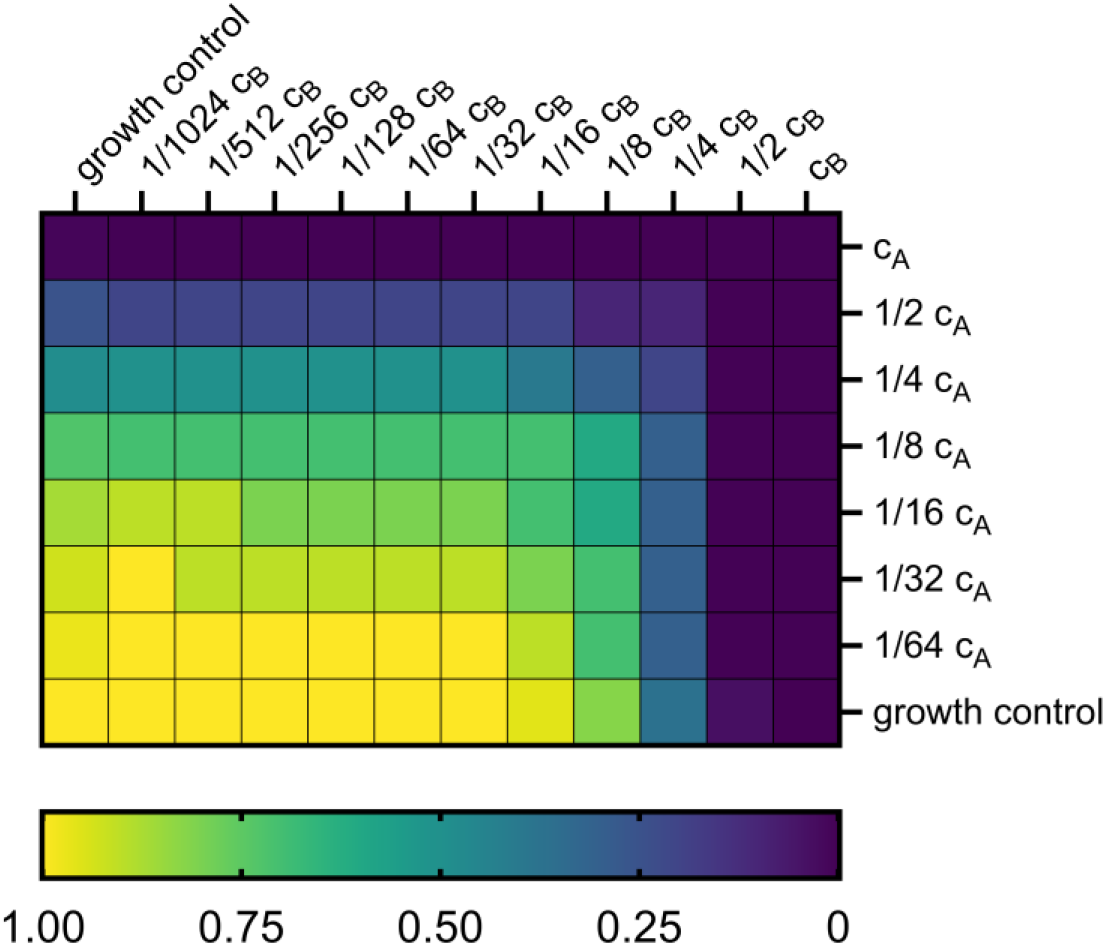
Heat map to demonstrate cell growth according to the concentration gradient with two serial diluted substances A (vertically) and B (horizontally). All values were normalized to growth control (lower left well).

The classification system proposed by Chou was used to classify the ∑FIC values determined experimentally according to equation (1) [17]. In this system, values ranging from 0.90 to 1.10 are classified as (nearly) additive effects, while values below 0.90 are described as indicating synergistic effects and values above 1.10 as indicating antagonistic effects. It was hypothesised that the closer the values are to 0, the stronger the synergistic behaviour. Consequently, the greater the difference from the value of 1.1, the more pronounced the antagonistic behaviour becomes, with no specified upper limit.

The investigation of the interactions between two hop isolates yielded different results depending on the bacterial strain.

**Fig.*5*** shows that additive effects could be detected for *B. subtilis* around a ∑FIC value of 1. Both the scattered dot plot and, in particular, the isobolograms showed that two hop compounds do not influence each other regarding their antibacterial activity. On the other hand, the investigation of *M. luteus* (**Fig.*6***) showed that the combination of two substances can lead to more than just additive effects. While the combination of humulone + lupulone (H + L) and humulone + isohumulone (H + iH) showed ∑FIC values below 1 and thus interact synergistically, the combination of isohumulone (iH) with xanthohumol (X) or isoxanthohumol (iX) resulted in antagonistic effects. Again, the plotted isobolograms support these findings.

**Fig. 5.**
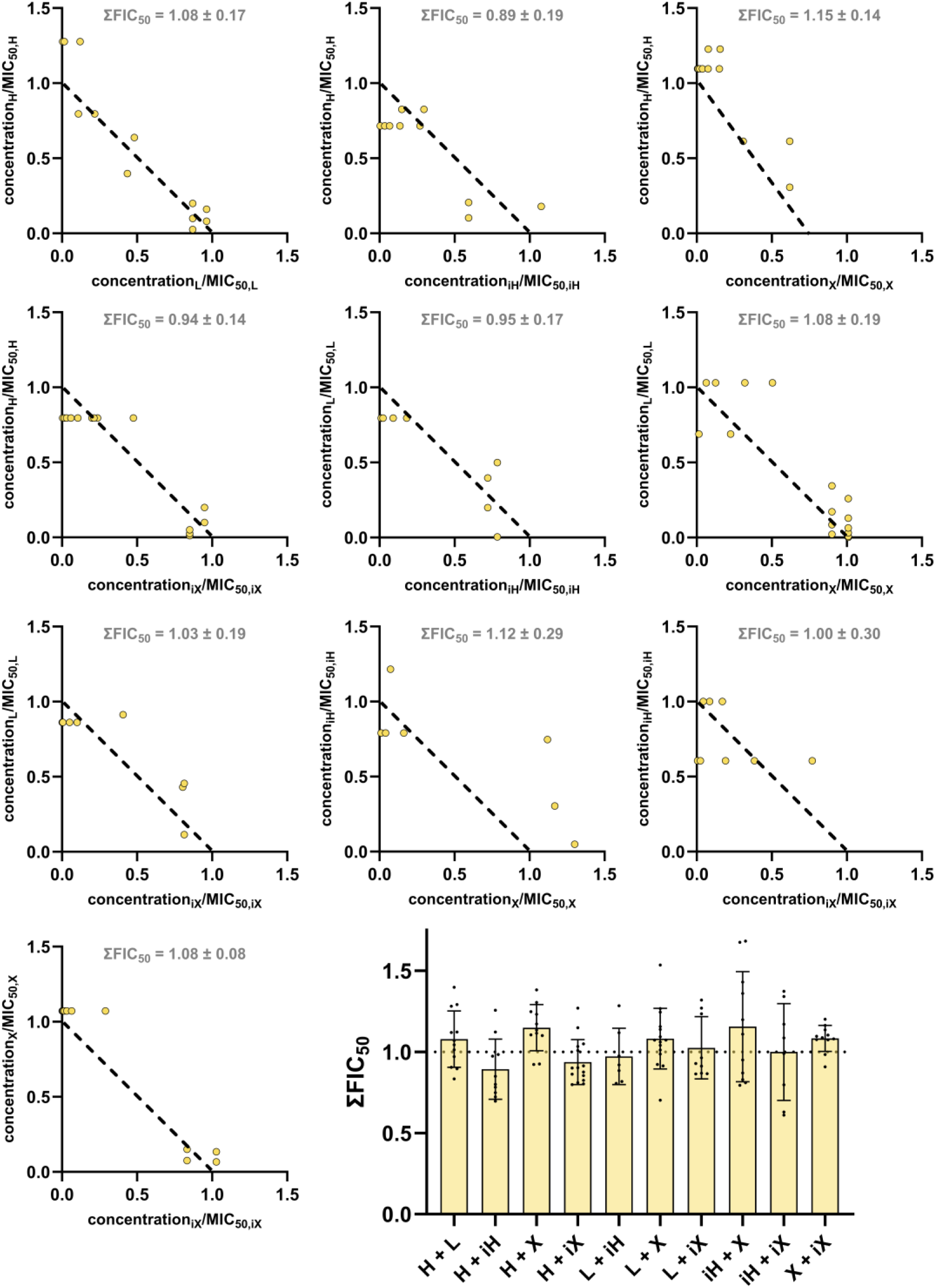
Isobolograms for all ten drug combinations possible against *B. subtilis*. Abbreviations refer to the hop isolates and therefore represent humulone (H), lupulone (L), isohumulone (iH), xanthohumol (X) and isoxanthohumol (iX). The distribution of data points is displayed using a scattered dot plot. Means and standard deviations were generated from at least 8 data points from two different plates.

**Fig. 6.**
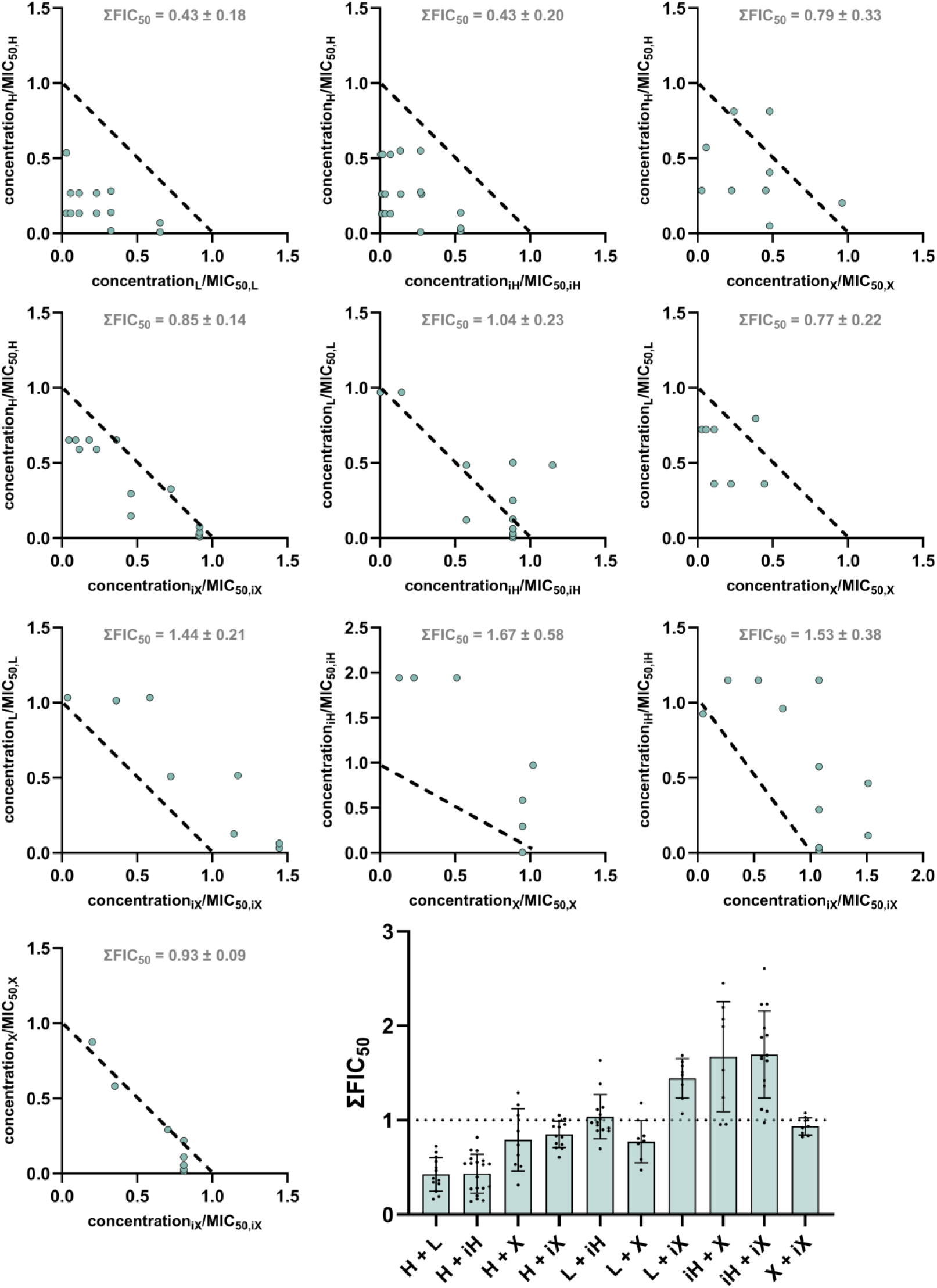
Isobolograms for all ten drug combinations possible against *M. luteus*. Abbreviations refer to the hop isolates and therefore represent humulone (H), lupulone (L), isohumulone (iH), xanthohumol (X) and isoxanthohumol (iX). The distribution of data points is displayed using a scattered dot plot. Means and standard deviations were generated from at least 8 data points from two different plates.

### 2.3. Cytotoxic correlation patterns on UMNSAH/DF-1

However, not only the antibacterial activity should be considered, but also the cytotoxic activity on the chicken cell line UMNSAH/DF-1 selected here. The interaction effects observed in **Fig.*7***are rather additive in their nature with a tendency toward antagonistic behaviour, which is favourable in the case of cytotoxic activity on a healthy chicken cell line. Only the combination of isohumulone and isoxanthohumol (iH + iX) led to synergistic effects with a ∑FIC value of 0.50 ± 0.23.

**Fig. 7.**
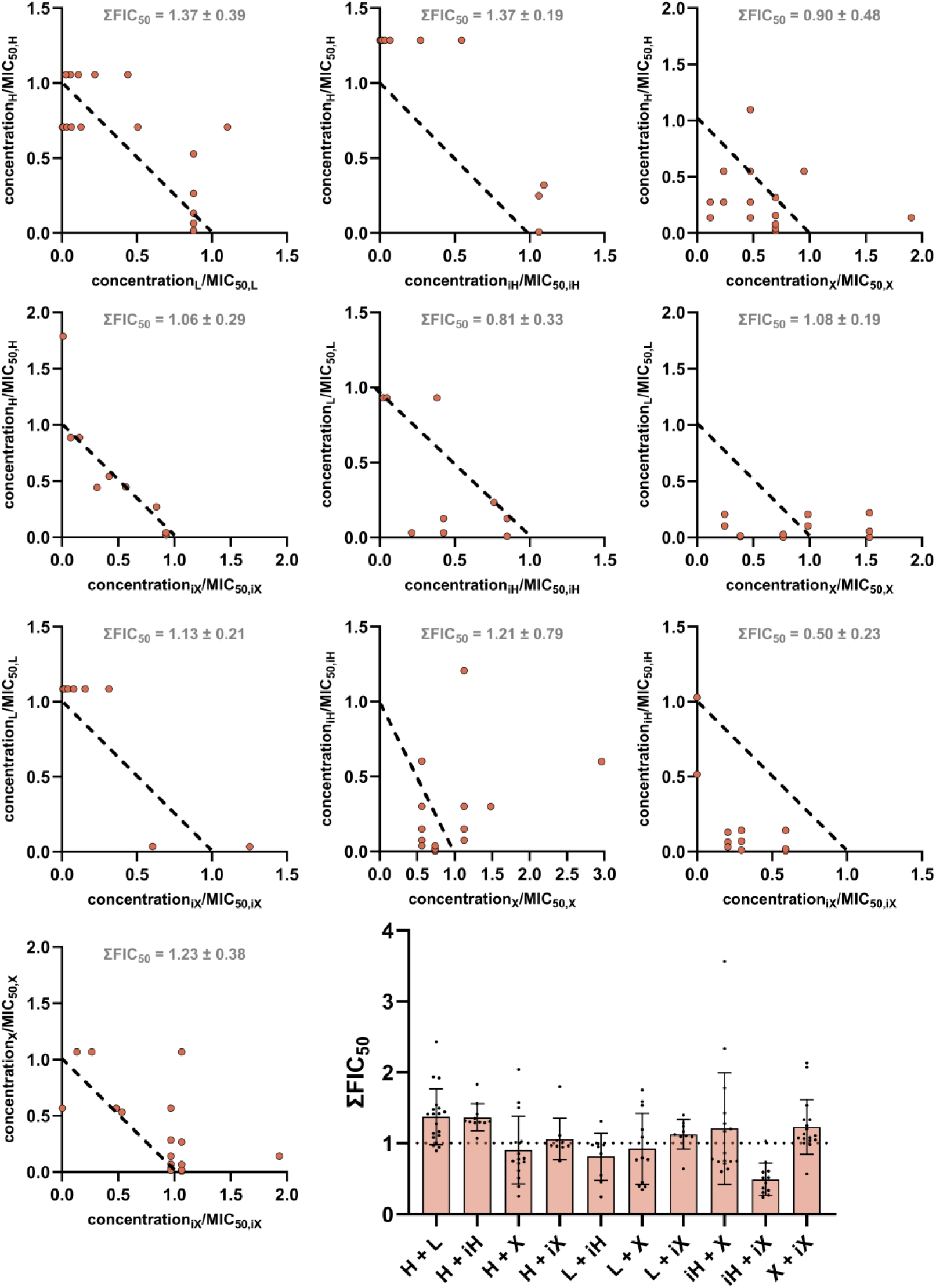
Isobolograms for all ten drug combinations possible against UMNSAH/DF-1. Abbreviations refer to the hop isolates and therefore represent humulone (H), lupulone (L), isohumulone (iH), xanthohumol (X) and isoxanthohumol (iX). The distribution of data points is displayed using a scattered dot plot. Means and standard deviations were generated from at least 8 data points from two different plates.

### 2.4. Selectivity of hop isolates with regard to their correlation patterns

In general, the highest possible antibacterial activity and the lowest possible cytotoxicity are beneficial for the use as a PFA. For easy evaluation, selectivity indices (SI) can be calculated by dividing the cytotoxic ∑FIC_50_ by the antibacterial ∑FIC_50_ for the respective pair of hop isolates. A value above 1 is desirable, as this indicates that the antibacterial effect is greater than the cytotoxic effect.

**Fig.*8*** shows the calculated selectivity indices. Once again, differences between the two bacterial strains can be seen here. While the selectivity indices for *B. subtilis* remain within the range of 1, higher indices can be observed for *M. luteus*. In general, this graph illustrates that, regardless of the strain selected, the combination of humulone and lupulone, as well as humulone and isohumulone, appears to be most favorable, whereas the combination of isohumulone and isoxanthohumol had the least selective effect.

**Fig. 8.**
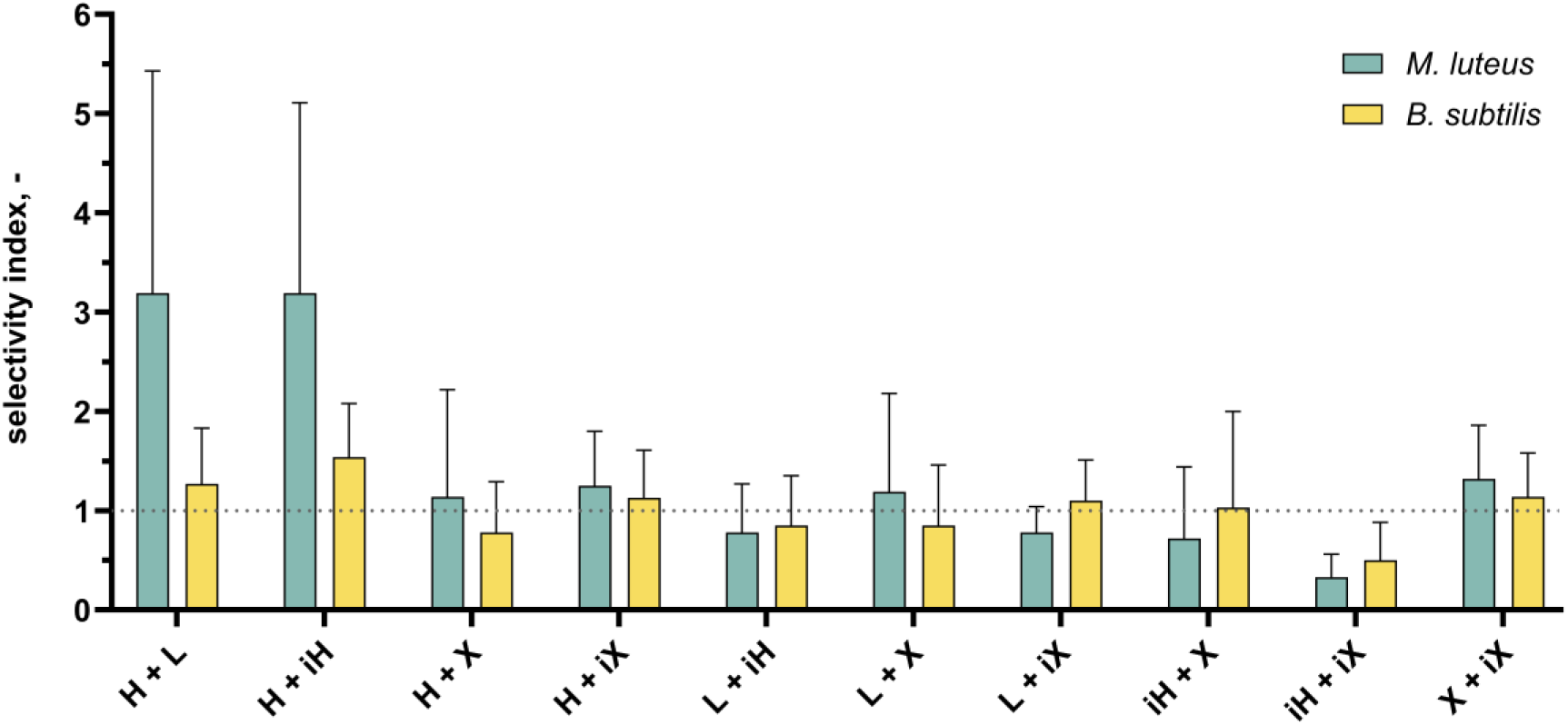
Selectivity for different hop isolate combinations on their antibacterial vs. cytotoxic activities. Abbreviations refer to the hop isolates and therefore represent humulone (H), lupulone (L), isohumulone (iH), xanthohumol (X) and isoxanthohumol (iX).

## 3. DISCUSSION

In general, ∑FIC values tend to have high standard deviations, which are often due to nonlinear dose-response relationships, exacerbated by the additional nonlinearity of drug interactions. Thus, when active ingredients have synergistic or antagonistic effects, even small fluctuations in concentration can lead to disproportionately large changes in MIC_50_ or IC_50_ values, thereby increasing the variability in FIC calculations. For the hop isolates, the scattering is additionally increased by the fact that the respective MIC_50_/IC_50_ values differ from one hop isolate to another. As a consequence, ΣFIC values may appear unstable or scattered as well. Nevertheless, trends for the correlation effects of hop isolates could be clearly identified, which are represented by underlying scattered dot plots and isobolograms.

### 3.1. Antibacterial correlation effects depend strongly on used bacterial strain

While the correlation between the antibacterial activities of hops and common antibiotics has already been researched in the literature [18], little is known about the potency of hop isolates themselves. Also, for application as PFA, the use in combination with conventional antibiotics is not effective, as these have been banned for prophylactic administration in many countries [3].

**Fig.6** and **Fig.5** showed that the combination of two hop isolates leads to additive relationships on *B. subtilis*, while all cases of correlation (synergism, additivity, and antagonism) occurred for *M. luteus. B. subtilis* is known for its elevated metabolic flexibility and the presence of alternative signalling pathways. Furthermore, the general stress response is more prominent in this case. In summary, substances can exhibit a more robust buffering effect in this instance, which frequently results in the occurrence of additive effects [19]. Conversely, *M. luteus* exhibits a reduced level of complexity, possessing a more limited repertoire of transporters and regulators, which therefore results in a weaker stress response. Consequently, substances can exhibit enhanced intermolecular interactions, resulting in non-linear effects [20]. In this study, only the combinations of lupulone and isohumulone (L + iH), as well as xanthohumol and isoxanthohumol (X + iX), showed rather linear effects, which in the case of L + iH could be attributed to relatively wide scattering or, in the case of X + iX, to the similarity of the structure. The most preferable synergistic effect on *M. luteus* showed the combination between humulone and lupulone (H + L) or humulone and isohumulone (H + iH), followed by humulone and xanthohumol (H + X), as well as lupulone and xanthohumol (L + X), indicating that combination of not isomerized forms of hop isolates are to be positively highlighted here.

### 3.2. Combination of hop isolate isoforms show synergistic cytotoxic behavior

The predominantly additive effects observed in healthy chicken cell line UMNSAH/DF-1 may be attributed to their intact regulatory and compensatory mechanisms, which allow effective buffering of multiple stressors. The cells can maintain metabolic and signalling homeostasis through coordinated stress responses and redundant pathways, preventing non-linear amplification of combined effects [21]. Consequently, interactions between compounds are more likely to sum up rather than showing synergistic or antagonistic actions. The only notable exception was the combination of isohumulone and isoxanthohumol, which ultimately led to a ∑FIC value clearly under 1 and therefore showed synergistic responses. While the underlying mechanism remains to be clarified, this observation underscores the potential benefits of avoiding both isoforms in the subsequent PFA. This objective could be realised with relative ease by utilising fresh hops or hop pellets and not waste streams of brewing processes.

### 3.3. Balancing antibacterial activity and cellular toxicity

In order to better understand the aforementioned effects in context, selectivity indices were calculated, which ideally highlights certain combinations of hop isolates more clearly. The selectivity indices concisely illustrate that the combination of humulone and lupulone should be particularly favoured for PFA applications. Humulone with its isomerized form isohumulone also achieved similar selectivity indices, but isohumulone performed rather poorly in combination with isoxanthohumol, which was the only combination to achieve an index of less than 1 for both bacterial strains *B. subtilis* and *M. luteus*. This also supports the hypothesis that fresh hops or hop pellets containing humulone, lupulone, and xanthohumol, but not their heat-formed isomers, can be used for the production of PFA.

Our findings might apply also to chicken pathogens as well, which are relevant targets for phytogenic feed additives. Chicken pathogens, such as Gram-positive *Clostridium perfringens* [22] exhibit intermediate genomic and metabolic complexity and are therefore more versatile than simple organisms like *Micrococcus luteus*, but less so than model bacteria like *Bacillus subtilis*. This suggests that insights into bacterial responses relevant to infection could be inferred using non-pathogenic model strains.

## 4. CONCLUSIONS AND OUTLOOK

This study provides a detailed analysis of the interaction patterns of hop isolates in both bacterial and eukaryotic cell systems, highlighting how these effects depend strongly on the complexity and metabolic characteristics of the target organism. On *Bacillus subtilis*, combinations of hop isolates generally resulted in additive effects, reflecting the bacterium’s high metabolic flexibility, alternative signaling pathways, and robust general stress response. In contrast, *Micrococcus luteus*, which possesses a simpler regulatory network and fewer transport systems, exhibited more variable responses, including synergistic and antagonistic interactions. These observations emphasize that the physiological complexity of the target organism strongly shapes the outcome of compound combinations. Among the tested combinations, humulone paired with lupulone consistently demonstrated favorable antibacterial activity while maintaining low cytotoxicity in UMNSAH/DF-1 chicken cell line with a selectivity index around 3. Notably, the combination involving isoforms isohumulone and isoxanthohumol resulted in synergistic cytotoxicity, suggesting that the use of fresh hops, containing the non-isomerized forms, may be preferable for the production of phytogenic feed additives. Crucially, these findings might be extended to chicken pathogens of agricultural relevance, such as *Clostridium perfringens* in future studies.

Overall, these results highlight that careful selection of hop isolate combinations can maximize antibacterial efficacy while minimizing host cytotoxicity. Such an approach provides a rational basis for the development of PFAs that are both effective and safe, with potential applications in poultry health management. The study underscores novel insights into the importance of considering target organism complexity, compound characteristics, and combination effects in the design of biogenic additive strategies. Future studies should focus on utilizing whole hop products instead of hop isolates to gain even more knowledge about compound interaction *in vitro* and their further application *in vivo*.

## 5. MATERIALS AND METHODS

### 5.1. Bacterial Strain, Media and Culture Conditions

*Micrococcus luteus* (DSM 1605) was obtained from DSMZ (German Collection of Microorganisms and Cell cultures GmbH). It was cultivated on nutrient agar or in nutrient broth (NB, DSMZ #1) containing 5 g L^-1^ peptone from casein (Carl Roth, Karlsruhe, GER) and 3 g L^-1^ meat extract (Carl Roth, Karlsruhe, GER) under aerobic conditions at 37 °C.

*Bacillus subtilis* (DSM 23778) was obtained from DSMZ (German Collection of Microorganisms and Cell cultures GmbH). It was cultivated on TSB (tryptic soy broth, DSMZ #545) agar plates or in TSB containing 17 g L^-1^ peptone from casein, 3 g L^-1^ peptone from soy, 2.5 g L^-1^ D (+) glucose, 5 g L^-1^ NaCl and 2.5 g L^-1^ K_2_HPO_4_ (Carl Roth, Karlsruhe, GER) under aerobic conditions at 28 °C.

For preparation of precultures, 20 mL of the respective medium was inoculated with a colony from a freshly streaked agar plate in 100 mL shaking flasks. Precultures were then incubated for 16 hours at either 37 °C for *M. luteus* or 30 °C for *B. subtilis*, with shaking at 180 rpm and a 50 mm shaking throw (Infors-HT, Bottmingen, Switzerland).

### 5.2. Chicken Cell Line, Media and Culture Conditions

UMNSAH/DF-1 cell line was purchased from Cytion (Eppelheim, GER). The cells were maintained in DMEM (Dulbeccos Modified Eagle Medium, Sigma-Aldrich, St. Louis, USA) containing 10% heat-inactivated FBS (fetal bovine serum, Sigma-Aldrich, St. Louis, USA) and cultivated at 37 °C and 5% CO_2_ in a humidified atmosphere. Cell lines were serial passaged after being detached with Accutase® solution (Sigma-Aldrich, St. Louis, USA).

### 5.3. Chromatographic Analysis of Hop Compounds

High performance liquid chromatography (HPLC) measurements were performed using a NUCLEODUR C18 Gravity-SB (3 µm, 250 x 2 mm) column equipped with a precolumn (Machery Nagel, Düren, GER) in a Shimadzu HPLC system (Kyoto, Japan) as described in a previous study [15].

### 5.4. Hop Isolates Preparation in Aqueous Media

The preparation of hop compound solutions was carried out analogously for NB, TSB and DMEM media for use in bacterial and avian cell culture. Hop compounds were provided by Hopsteiner (Mainburg, GER) in the following purities: isoxanthohumol (81.0%), xanthohumol (90.0%), humulone (88.5%), lupulone (74.3%) and isohumulone (91.0%). The hop compounds were dissolved in ethanol to 10 mg mL^-1^. From the stock solution, 1:10 dilutions were prepared in the respective medium, resulting in precipitation due to their hydrophobic character. In addition, 1:10 dilutions of the stock solution were carried out in bacterial medium containing 10 mM (2-hydroxypropyl)-β-cyclodextrin (CD, Sigma-Aldrich, St. Louis, USA) in order to increase the solubility of the hydrophobic compounds. Both approaches (with and without CD) were followed by centrifugation (10.000 xg, 10 min). After that, the clear supernatant was transferred to a new tube and used for further experiments. Due to the low solubility of the compounds, the actual dissolved concentration was in most cases lower than the weighed concentration. This is why the actual concentration of the compounds in the aqueous medium was analysed using HPLC and used for subsequent calculations.

### 5.5. 2D Checkerboard assay for antibacterial correlation pattern

Checkerboard assay was performed based on Bellio et a*l*. [7]. First, 100 µL of the respective medium was pipetted into 96-well microtiter plates (Sarstedt, Nuembrecht, GER), before prepared hop isolates were diluted vertically and horizontally according to method description. This was done for each combination of compounds, resulting in a total of ten different combinations. Two plates were used for each combination, with the layout of the substances being reversed on one of the plates. After that, the optical density (OD) of a preculture prepared on the previous day was measured at 600 nm in a spectrophotometer (Implen, Muenchen, GER). The inoculum was then diluted with respective medium to a desired OD_600nm_ of 0.1 and transferred to all wells (100 µL). The background signal (OD_750nm_) was measured immediately afterwards using a plate reader (Tecan, Maennedorf, CH). The chosen wavelength of 750 nm resulted from previous experiments with hop extracts in order to avoid an overlap with the absorption spectrum of chlorophyll [23]. Final measurements were performed after 6 h for *B. subtilis* and 24 h for *M. luteus* based on their growth rates. Data analysis, which will be described later, was the same for antibacterial and cytotoxic effects.

### 5.6. 2D Checkerboard Assay for cytotoxic correlation pattern

In order to investigate possible correlations in the cytotoxic effect of the hop isolates, the checkerboard assay described above was adapted for the adherent UMNSAH/DF-1 cell line. Cells were seeded in 96-well microtiter plates (Sarstedt, Nuembrecht, GER) with a cell density of 0.05 x 10^6^ cells per mL and 100 µL per well and incubated for 24 h (37 °C, 5% CO_2_). Analogous to the previous descriptions, this time deep-well plates (Greiner Bio-One GmbH, Frickenhausen, GER) were prepared the next day using the dilution scheme. After that the dilution series was transferred to the cell-seeded plates (100 µL). In addition, plates without cells and only with medium were prepared in this way, which were used for the background measurements. Treated cells and background plates were further incubated for 48 h (37 °C, 5% CO_2_). The cytotoxic effect was analysed using the MTT (3-(4,5-dimehtylthiazol-2-yl)-2,5-diphenyltetrazolium bromide) based cell viability assay. For that 12.5 µL of MTT (Sigma-Aldrich, St. Louis, USA) solution (0.5% (w/v) in phosphate-buffered saline (PBS)) was added per well, followed by another 3 h of incubation (37 °C, 5% CO_2_). The plates were centrifuged (300 xg, 5 min), the supernatant was discarded and 20 µL of Igepal (0.4% (v/v) in H_2_O, Sigma-Aldrich, St. Louis, USA) was added. After incubation on a plate shaker for 10 min at 1.000 rpm, 100 µL per well of DMSO was added to dissolve the formazan. The plates were incubated again on the plate shaker for 30 min. Absorbance at 570 nm was measured using a plate reader (PerkinElmer, Waltham, USA). The resulting absorbance was directly proportional to the amount of viable cells.

### 5.7. Data analysis

For data analysis, the background signal was subtracted. For bacterial approaches, the background signal was measured immediately after addition of the bacterial suspension, whereas for cytotoxic approaches MTT-treated compound plates without cells served as background signal. All values were normalized to the growth control (lower left well). Subsequently, the MIC_50_/IC_50_ was determined for each individual substance, based on the rows in which only one substance was diluted. MIC_50_ (minimum concentration of a substance that inhibits the growth rate of the bacteria tested by 50%) and IC_50_ (concentration of a substance that inhibits the metabolic activity of the cell line tested by 50 %) values were calculated by non-linear regression using the GraphPad Prism model ‘log(inhibitor) vs. response’, where the bottom was set to a constant value (0). The fractional inhibitory concentration (∑FIC) was estimated for all wells containing both compounds in combination and exhibiting a value of 0.5 after normalisation to the growth control. ∑FICs were calculated according to general equation of Loewe additivity (Eq. (1)), where d_1_ and d_2_ are doses of each drug in combination that yield the same effect as D_1_ or D_2_ (1) given alone [8]. The effect observed in this study is the half-maximal growth rate inhibition of *M. luteus* and *B. subtilis* or half-maximal inhibition of metabolic activity of UMNSAH/DF-1.

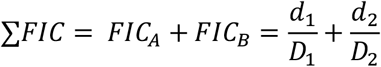

Means and standard deviations (SDs) were calculated from two plates with at least eight data points. In addition, isobolograms were also created for visualisation purposes and better evaluation of the effects.

## 7.ACKNOWLEDGEMENTS

This study was funded by the German Bundesministerium für Forschung, Technologie und Raumfahrt (BMFTR, Grant-No. 031B1253). Hop compounds were kindly provided by Hopsteiner (Hallertauer Hopfenveredelungsgesellschaft mbH, Mainburg, DE).

## 8. AUTHOR CONTRIBUTIONS

**Luisa Kober:** Conceptualization (equal); writing – original draft (lead); experiments and analysis (lead); writing – review and editing (equal)

**Luca von Karger:** Experiments and analysis (supporting); writing – review and editing (supporting)

**Kathrin Castiglione:** Conceptualization (equal); writing – original draft (supporting); writing – review and editing (equal)

## 9. DATA AVAILABILITY

All data generated and analysed during this study are included in this published article. Further data is available from the corresponding author on reasonable request.

## 10. ADDITIONAL INFORMATION

### 10.1. ETHICS APPROVAL AND CONSENT TO PARTICIPATE

Not applicable.

### 10.2. CONSENT FOR PUBLICATION

Not applicable.

